# INRI-seq enables global cell-free analysis of translation initiation and off-target effects of antisense inhibitors

**DOI:** 10.1101/2022.04.11.487859

**Authors:** Jens Hör, Jakob Jung, Svetlana Đurica-Mitić, Lars Barquist, Jörg Vogel

## Abstract

Ribosome profiling (Ribo-seq) is a powerful method for the transcriptome-wide assessment of protein synthesis rates and the study of translational control mechanisms. Yet, Ribo-seq also has limitations. These include difficulties with detection of low abundance transcripts and analysis of translation-modulating molecules such as antibiotics, which are often toxic or challenging to deliver into living cells. Here, we have developed *in vitro* Ribo-seq (INRI-seq), a cell-free method to analyze the translational landscape of a fully customizable synthetic transcriptome. Using *Escherichia coli* as an example, we show how INRI-seq can be used to analyze the translation initiation sites of a transcriptome of interest. We also study the global impact of direct translation inhibition by antisense peptide nucleic acid (PNA) to analyze PNA off-target effects. Overall, INRI-seq presents a scalable, sensitive method to study translation initiation in a transcriptome-wide manner without the potentially confounding effects of extracting ribosomes from living cells.

## INTRODUCTION

Protein synthesis is one of the most energy-consuming processes in living cells, making its precise regulation a crucial matter of cellular economy^1, 2^. While mRNA levels determined by RNA-sequencing (RNA-seq) are often used as a proxy for protein synthesis in global gene expression analysis^3^, final protein abundance does not always correlate with mRNA levels^4, 5, 6, 7^. This is due to a multitude of regulatory mechanisms, for example, direct control of the translational machinery^1^ and post-transcriptional control of mRNAs by base pairing small RNAs (sRNAs)^8^ or intrinsic mRNA structure^9^.

Over the past decade, ribosome profiling (Ribo-seq) has become a primary method to more directly measure protein synthesis in a transcriptome-wide manner^10, 11, 12^. Ribo-seq is based on RNA-seq analysis of ribosome-protected fragments (RPFs), which are ribosome-bound mRNA fragments that survive nuclease treatment after cell lysis because they are covered and protected by translating ribosomes^13, 14, 15^. In *Escherichia coli*, RPFs typically are 15 to 45 nt in length^16^. Since each RPF represents the position of one ribosome, their sum not only reveals which mRNAs but also which parts of the coding sequence (CDS) of these mRNAs were being translated at the point of sampling. Thus, Ribo-seq can map the landscape of actively translating ribosomes *in vivo* to provide global information about translational pausing^16^, stalling^17^ and start site usage^18, 19, 20^, as well as an estimate of protein copy numbers^6^.

While Ribo-seq has greatly advanced the study of translation-related processes, the method has not been without limitations. Coverage of weakly expressed genes remains challenging, preventing many genes from being detected in common study designs. This includes certain gene classes with notoriously low expression under standard growth conditions, such as toxins whose expression is only triggered by specific stresses that lead to cell death or inhibition of growth^21^. Similarly, Ribo-seq of microbes from important ecological habitats such as the human gut^22^ remains difficult since many of them cannot be cultured in the laboratory, though recent efforts have started to tackle this challenge^7^.

On the mechanistic level, Ribo-seq-based studies of molecules affecting translation can be hampered by cellular responses. For instance, the antibiotic retapamulin (RET) acts right after translation initiation when the ribosome is still at the start codon, which has enabled the global annotation of translation initiation sites (TISs) in *E. coli*^18, 23^. However, this study required genetic inactivation of the ABC transporter TolC to prevent export of the antibiotic prior to it exerting its effect. While this was possible in *E. coli*, a lack of either genetic tools or knowledge of transporters would preclude similar studies in many other organisms. Lastly, since Ribo-seq is performed on living cells, it can be difficult to dissect direct and indirect effects on translation. This is exemplified by antisense antibiotics^24, 25, 26^, whose import via carrier peptides broadly affects gene expression in addition to the desired antisense inhibition of the targeted gene of interest^27, 28^.

To overcome some of these limitations, we have developed *in vitro* Ribo-seq (INRI-seq) for the global study of translation in a cell-free manner. INRI-seq uses the commercially available PURExpress *in vitro* translation system combined with an *in vitro*-synthesized, fully customizable transcriptome for better control of individual mRNA levels. INRI-seq obviates the need for translation-modulating compounds to traverse cellular membranes and the extraction of ribosomes from a large number of living cells. As proof of concept, we apply INRI-seq to a synthetic *E. coli* transcriptome and show that the method faithfully validates known TISs and can be used to predict new TISs. In addition, we use the system to study the fidelity of translational inhibition by antisense peptide nucleic acid (PNA), demonstrating on-target specificity and defining base-pairing criteria that influence the off-target effects of these short antisense oligomers. INRI-seq bears great potential as a scalable alternate method to study translation control mechanisms and translation-modulating compounds in other organisms and organismal communities, including eukaryotes.

## RESULTS

### Design and workflow of INRI-seq

To obtain a synthetic transcriptome for INRI-seq, we designed a single-stranded DNA oligonucleotide pool consisting of 4,386 unique oligonucleotides covering all annotated CDSs of *E. coli* K-12 MG1655 (NC_000913.3). For each CDS, we included the natural, i.e. annotated, start codon (generally ATG, GTG or TTG) followed by the first 150 nt of the open reading frame (ORF) (Figure 1). We also included the ribosome binding site by addition of 30 nt of the respective 5′ untranslated region (UTR), corresponding to the average length of 5′ UTRs in *E. coli*^29^. At the 3′ end, we added a strong TAA stop codon^30^ and the 3′ UTR of the *E. coli hns* gene for uniform termination of translation. Importantly, this 3′ UTR doubled as primer binding site to generate a double-stranded DNA (dsDNA) pool. Finally, a T7 RNA polymerase promoter was added to the 5′ end of each oligonucleotide to facilitate *in vitro* transcription, while at the same time representing the second primer binding site for pool amplification. The sequence in between the flanking regions (T7 promoter and 3′ UTR) is completely flexible, and could be replaced with the sequence of any gene of interest.

**Figure 1.**
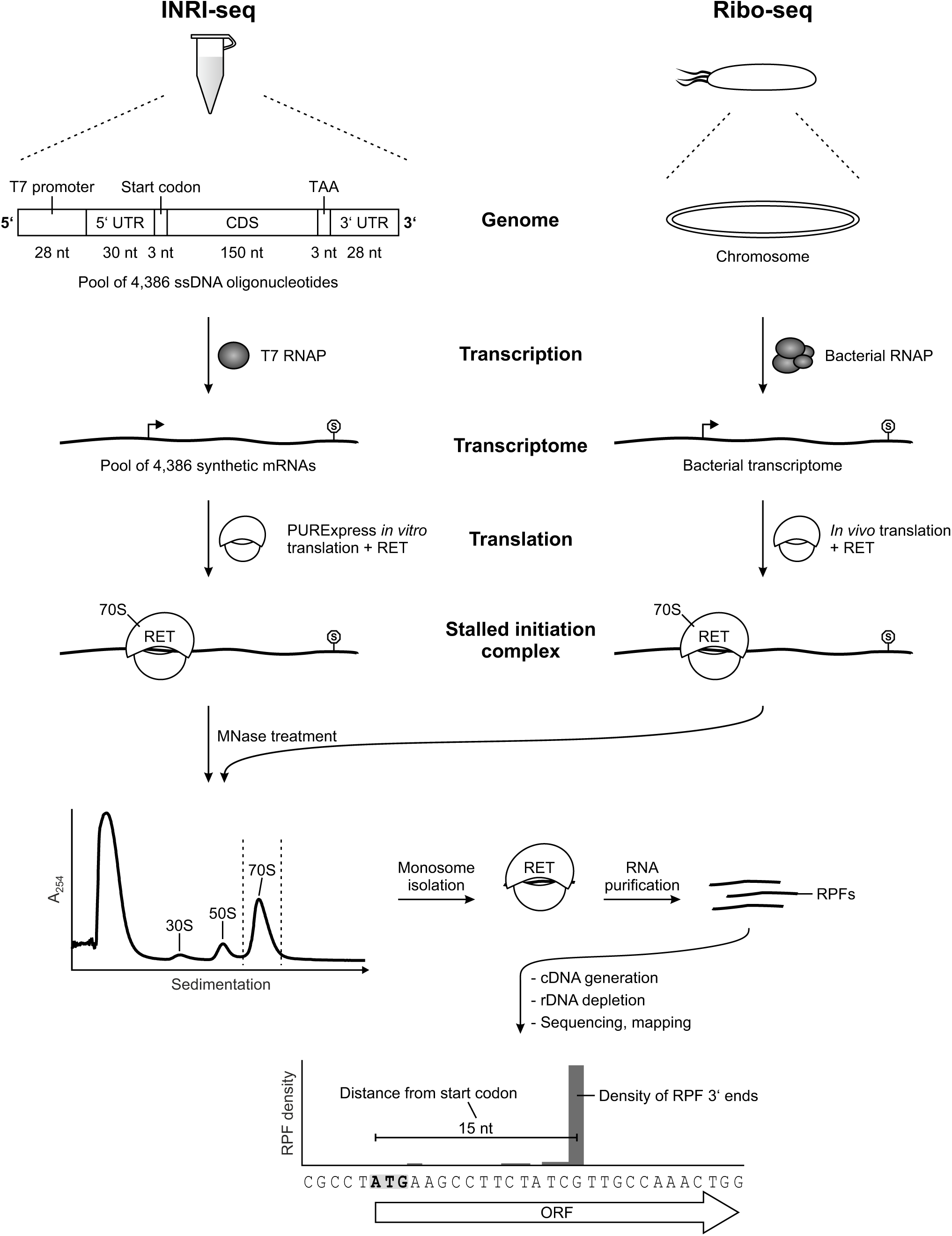
Workflow of INRI-seq. INRI-seq employs a synthetic transcriptome, for the generation of which one ssDNA oligonucleotide is synthesized for each gene of *E. coli* MG1655. Every gene contains a T7 RNA polymerase promoter, 30 nt of its natural 5′ UTR, its natural start codon, the first 150 nt of its CDS, a TAA stop codon, and a 28-nt 3′ UTR. This pool of oligonucleotides is converted to dsDNA by limited cycle PCR and then transcribed *in vitro* to obtain the synthetic transcriptome. Following *in vitro* translation of the synthetic transcriptome, RET is added and the translation continued, leading to ribosomes stalled at the start codons of the transcripts. RNA not protected by ribosomes is cleaved with MNase and the reaction is sedimented on a sucrose gradient. After 70S monosome isolation, RPFs are extracted from the ribosomes and size-selected on a gel. The purified RPFs are converted to cDNA, rDNA is depleted and the resulting sample is sequenced. Attributing the read density to the 3′ ends of the sequenced reads allows identification of the ribosome positions and with it analysis of TISs.

Following a limited-cycle PCR of the oligonucleotide pool, we used the resulting dsDNA as template for T7 *in vitro* transcription. RNA-seq analysis of the obtained synthetic transcriptome detected 4,225 of the 4,386 synthetic mRNAs at an abundance of >10 reads per million (median abundance: 125 reads per million), showing almost complete (∼96%) coverage of the ORFs included in the original DNA pool design (Figure S1A).

Next, we applied the INRI-seq protocol to study the TISs of the synthetic *E. coli* transcriptome *in vitro*. We used RET, an antibiotic known to stall the ribosome at the start codon immediately after initiation (Figure 1)^18, 23^. Since an initiating ribosome protects 14-16 nt of the mRNA downstream of the first nucleotide of the start codon^31^, the 3′ ends of RPFs will identify the TIS of the translated CDS following Ribo-seq^20^. Since RET does not inhibit translation elongation^18^, its addition to translating ribosomes causes polysomes to collapse into monosomes if translation is continued. We found that under our experimental conditions, addition of RET at a final concentration of 5 µg/ml for 30 min after initial translation for 15 min causes polysome collapse (Figure S1B). This is considerably longer than the 5 min of RET treatment necessary *in vivo*^18^, suggesting that *in vitro* translation of the shorter CDSs used here might be less efficient.

To purify RET-stalled monosomes, the associated transcripts were cleaved with MNase and the digested samples were subsequently run on sucrose gradients, as is common for bacterial Ribo-seq (Figure 1)^16^. After collection of the 70S peak, RPFs were extracted with acidic phenol-chloroform followed by size selection (15-45 nt) of RNA in a denaturing polyacrylamide gel. These isolated RPFs can directly be used for library preparation and sequencing (Figure 1). Preliminary sequencing showed 24 fragments of ribosomal RNAs (rRNA; 9 fragments for 23S rRNA, 13 for 16S rRNA, and 2 for 5S rRNA) to be strongly enriched in this short RNA fraction (Figure S1C). In the final INRI-seq protocol, these 24 rRNA fragments are depleted from the final cDNA library using a Cas9-based DASH protocol^32^, which reduces rRNA reads from >90% to <1% (Figure S1D, Supplementary Table S1). This *in vitro* Ribo-seq protocol enables us to globally study translation initiation in a synthetic transcriptome of *E. coli*.

### INRI-seq identifies annotated TISs

We sequenced five independent INRI-seq libraries to study the TISs of *E. coli*, as described above. All libraries correlated well, indicating high reproducibility between experiments (Figure S2A). Of the 4,225 transcripts present in the transcriptome (Figure S1A), 4,149 were detected in all five replicates (Figure S2B). By assigning the read densities to the 3′ ends of the sequenced RPFs, we observed an average distance of 15 nt from the first nucleotide of the annotated start codons, in line with the expected ribosome protection (Figure 2A, Figure S2C). Crucially, this distribution was fully consistent with the peak distribution of sequenced RPFs in an *in vivo* study using a similar RET-based protocol (Ribo-RET)^18^. Manual inspection of the data further revealed that the position of RPF density generally overlapped very well between the *in vitro* and the *in vivo* data, as exemplified by the *cspE* transcript, where the only detected density was close to the start codon (Figure 2B). Indeed, in both the INRI-seq and Ribo-RET datasets, the distance of the RPF peak density to the *cspE* start codon was exactly 16 nt (Figure 2C). The overlap was also present for transcripts with unexpected density distributions such as *hfq*, for which two peaks 13 and 16 nt downstream of the start codon were detected by both methods (Figure S2D, E).

**Figure 2.**
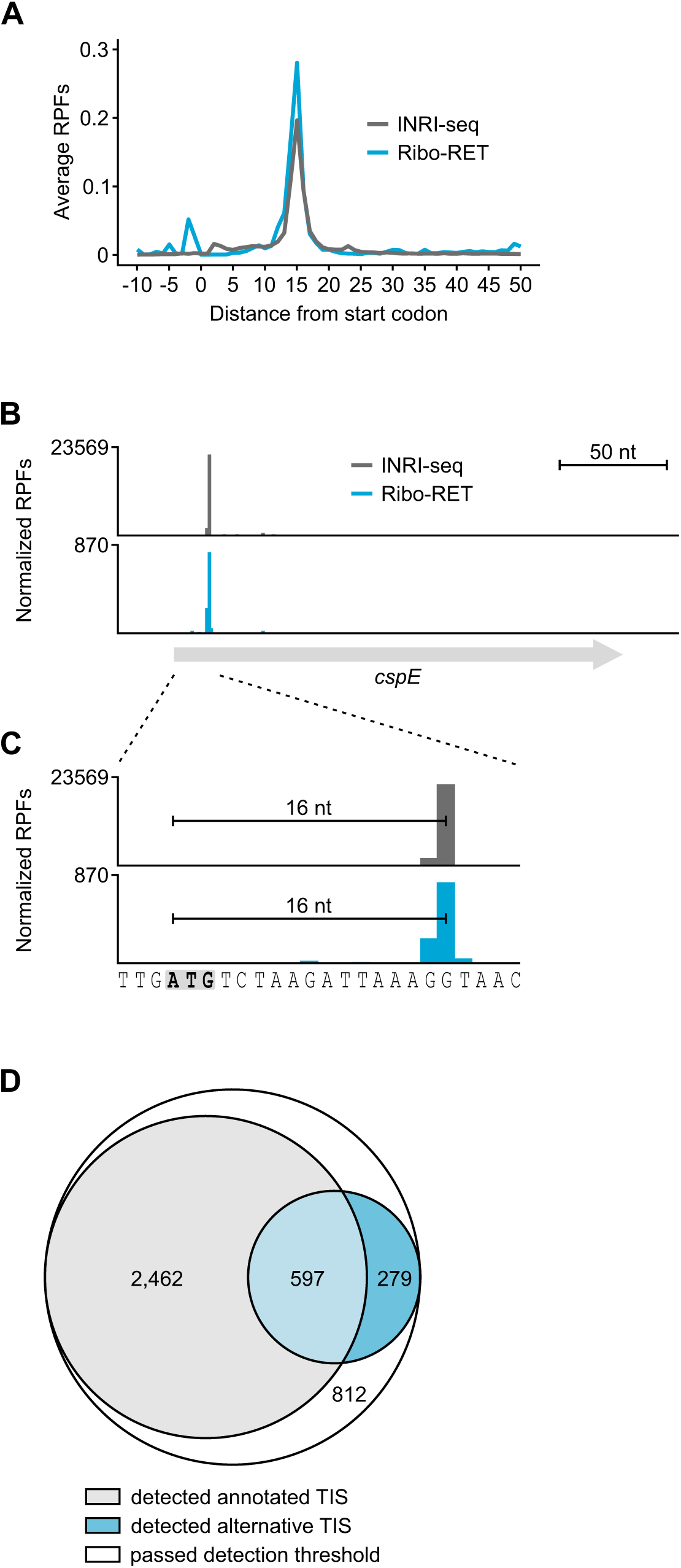
INRI-seq verifies annotated TISs. (A) The average distance of RPF peak density from annotated start codons is 15 nt, in agreement with previous *in vivo* Ribo-RET data^18^. (B) INRI-seq detects one RPF peak in the *cspE* gene. (C) INRI-seq RPF peak density is located exactly 16 nt downstream of the annotated start codon (gray and bold) agreeing with previous *in vivo* Ribo-RET data^18^. (D) Euler diagram showing the transcripts passing the thresholds for TIS detection. For 3,059 of these, a peak at the annotated start codon was detected, whereas one or more alternative TISs were detected for 858 of them.

Globally, INRI-seq confirmed the annotated TISs of 3,059 out of 4,386 (∼70%) genes using stringent cut-off criteria (RPF peak with counts per million mapped reads (CPM) ≥5 within 12-18 nt of the start codon in at least two of the five replicates; Figure 2D, Supplementary Table S2). Using the same criteria, *in vivo* Ribo-RET verified only 780 (∼18%) annotated TISs^18^ (Supplementary Table S2), suggesting that INRI-seq is more sensitive, likely due to more even transcript abundance in the pool. No TIS was detected for 812 transcripts that passed our detection thresholds (Figure 2D). For 99 of them, a peak was detected in only one of the five replicates, whereas the others did not show a clear peak that could be attributed to a TIS (Supplementary Table S2).

### INRI-seq confirms alternative TISs

Stalling ribosomes at start codons using antibiotics is a powerful way of analyzing where on a transcript translation initiation takes place^18, 19, 20^. In addition to validating known TISs, these kinds of datasets can also be searched for new TISs, revealing alternative in-frame (Figure 3A) or out-of-frame TISs (Figure 3B) within annotated genes. To investigate whether INRI-seq is able to identify alternative TISs, we compared our data to *in vivo* Ribo-RET data^18^. Of the 64 alternative TISs identified *in vivo*, we detected 51 (Supplementary Table S3). Most of the alternative TISs that were missed from the INRI-seq data set were >150 nt downstream of the annotated start codons and therefore not part of our synthetic transcriptome.

**Figure 3.**
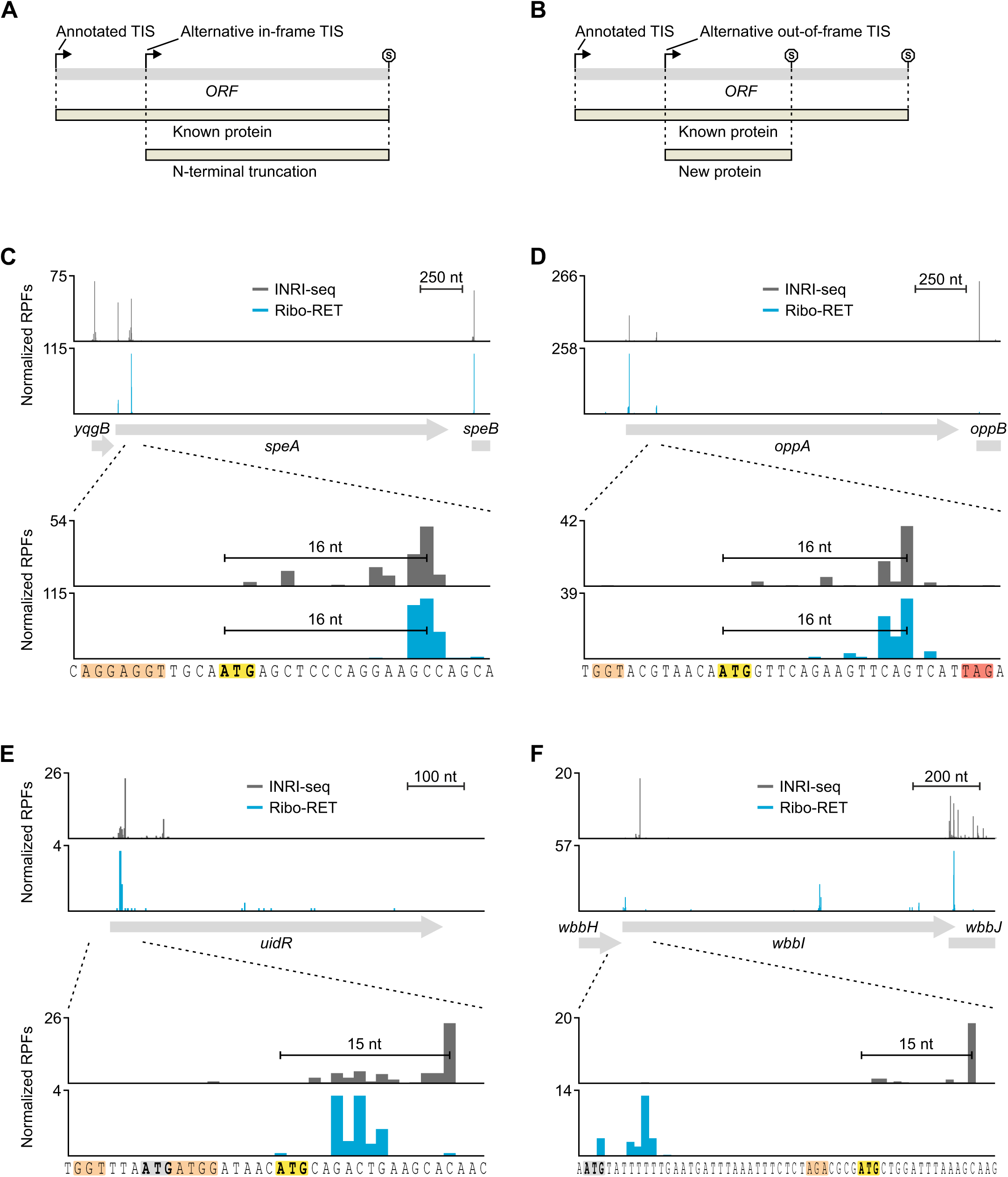
INRI-seq verifies known alternative TISs and identifies new putative TISs. (A) Alternative in-frame TISs lead to N-terminally truncated versions of known proteins. (B) Alternative out-of-frame TISs generate new proteins. (C) INRI-seq detects two RPF peaks at the 5′ end of *speA*. The second RPF peak identified in *speA* belongs to a known, in-frame alternative start codon^18^. (D) INRI-seq detects two RPF peaks at the 5′ end of *oppA*. The second RPF peak identified in *oppA* belongs to a known, out-of-frame alternative start codon^18^. (E) INRI-seq detects an RPF peak at the 5′ end of *uidR*. The detected RPF peak cannot be attributed to the annotated start codon of *uidR* but rather derives from an in-frame alternative start codon. (F) INRI-seq detects an RPF peak toward the 5′ end of *wbbI*. The detected RPF peak cannot be attributed to the annotated start codon of *wbbI* but rather derives from an out-of-frame alternative start codon. Gray and bold, annotated start codon. Yellow and bold, alternative start codon. Orange, SD sequence. Red, stop codon.

Next, we inspected some examples in more detail. The gene encoding arginine decarboxylase, *speA*, was previously shown to contain an in-frame alternative TIS that shortens the secretion signal-containing N-terminus of SpeA by 26 amino acids (aa) and leads to cytoplasmic rather than periplasmic localization of this isoform^18^. In accordance with the *in vivo* Ribo-RET data, INRI-seq displayed a strong peak of RPF density 16 nt downstream of this alternative start codon (Figure 3C). Importantly, INRI-seq also detected the start codon of the 43-aa short *yqgB* gene encoded upstream of *speA*, which was not possible *in vivo*, most likely due to its low abundance (Figure 3C). Different from the alternative in-frame TIS in *speA*, the *oppA* gene encoding an oligopeptide uptake protein as well as the OppX RNA sponge^33, 34^ contains an out-of-frame TIS for an alternative 7-aa ORF^18^. Our INRI-seq data supports this prediction, exhibiting the same RPF peaks as the *in vivo* data (Figure 3D). While it is unclear if this 7-aa peptide has a biological function, translation of this alternative frame could also impact the translation of *oppA*, similarly to upstream ORFs that can regulate translation of their downstream genes^35^. In conclusion, these examples illustrate how INRI-seq not only captures annotated TISs but also reports TISs that were not detected by *in vivo* Ribo-seq.

### INRI-seq identifies putative new TISs

After confirming that INRI-seq is able to detect both annotated and known alternative TISs with high sensitivity, we asked whether this method also enables the detection of putative new TISs. To do so, we searched for peaks with a relative density of ≥10% of the total reads identified for a given gene. These were further filtered to only include peaks of ≥5 CPM, which had to be present in at least two of the five replicates. Using these criteria, we identified 918 putative new TISs, 279 of which could be assigned to genes for which INRI-seq did not detect the annotated TIS (Figure 2D, Supplementary Table S3). We then studied two interesting candidates in more detail.

UidR is a transcriptional repressor of the *uid* operon, which is involved in transport and degradation of β-glucosides^36^. INRI-seq showed a clear RPF density peak toward the 5′ end of *uidR*, but downstream of the annotated start codon (Figure 3E). Instead, the distance of the identified peak agreed well with an in-frame AUG four codons downstream of the annotated start codon, suggesting that the UidR protein might be 4 aa shorter than annotated (Figure 3E). Indeed, this shorter N-terminus is supported by a recent mass spectrometry study of the N-termini of *E. coli* proteins, which identified the exact peptide (MQTEAQPTR) that corresponds to the TIS identified by INRI-seq^37^. By contrast, *in vivo* Ribo-RET^18^ showed only minimal density in this position, leaving the TIS of *uidR* ambiguous.

Another interesting alternative TIS candidate identified by INRI-seq is present in *wbbI* (Figure 3F), the gene encoding β-1,6-galactofuranosyltransferase^38^. As for *uidR*, no RPF density corresponding to the annotated start codon of *wbbI* was seen (Figure 3F). Instead, a peak corresponding to an out-of-frame AUG was detected close to the 5′ end of the gene. This internal ORF encodes for a 31-aa long peptide, for which we could not find any homologous domains or sequences using Pfam or PHMMER queries, respectively^39, 40^. Interestingly, although *in vivo* Ribo-RET did not identify this alternative TIS, it revealed another out-of-frame internal ORF encoding a 47-aa peptide within *wbbI*^18^. Due to its distance from the annotated start codon, INRI-seq was unable to detect this additional alternative TIS. Yet, it is intriguing to think that a single gene might contain two different out-of-frame ORFs.

Overall, INRI-seq is able to validate known alternative TISs and allows identification of putative new TISs, which should facilitate more detailed follow-up *in vivo* studies.

### Analysis of in vitro translation inhibition by PNAs

Translation inhibition by antisense oligomers, specifically peptide nucleic acid (PNA), is an upcoming field of antibiotics research. PNAs have the potential for species-specific killing, only targeting the pathogen while leaving beneficial microbiota untouched^24, 25, 26^. Similar to bacterial sRNAs^8^, antisense oligomers inhibit translation by blocking ribosome access to the ribosome binding site of their targets^41^. Yet, the high sample volumes required by standard Ribo-seq studies pose a challenge for studies of translation control and potential off-targeting by such antisense antibiotics *in vivo*, because of the high manufacturing costs of PNAs. We therefore decided to exploit the small-volume, cell-independent nature of INRI-seq to quantitatively study the effects of varying PNA concentrations on translation in a transcriptome-wide manner.

To test whether PNA-mediated translation inhibition can be analyzed *in vitro*, we adopted the well-established translation inhibition by a 10-mer antisense PNA that sequesters the start codon region of the *E. coli acpP* mRNA, which encodes the essential acyl carrier protein (Figure 4A)^28, 42, 43^. To this end, we created a translational fusion transcript containing 50 nt of the 5′ UTR and the first 8 codons of *acpP* fused to the CDS of *gfp*. Upon addition of increasing concentrations of *acpP*-PNA, *in vitro* synthesis of the AcpP::GFP fusion protein was strongly reduced (Figure 4B). Importantly, this was not the case when a scrambled version of the PNA (*acpP*-PNA-scr) was added to the reaction. At an equimolar ratio of *acpP*-PNA to *acpP*::*gfp* (100nM), translation was inhibited by ∼17%, whereas a 5-fold excess of *acpP*-PNA caused a ∼74% drop in translation (Figure 4C). These results are consistent with a previous *in vitro* study analyzing PNA efficiency in a less-defined reticulocyte extract using DNA as template^44^ and demonstrate that our *in vitro* system can be used to study the effect of antisense oligomers.

**Figure 4.**
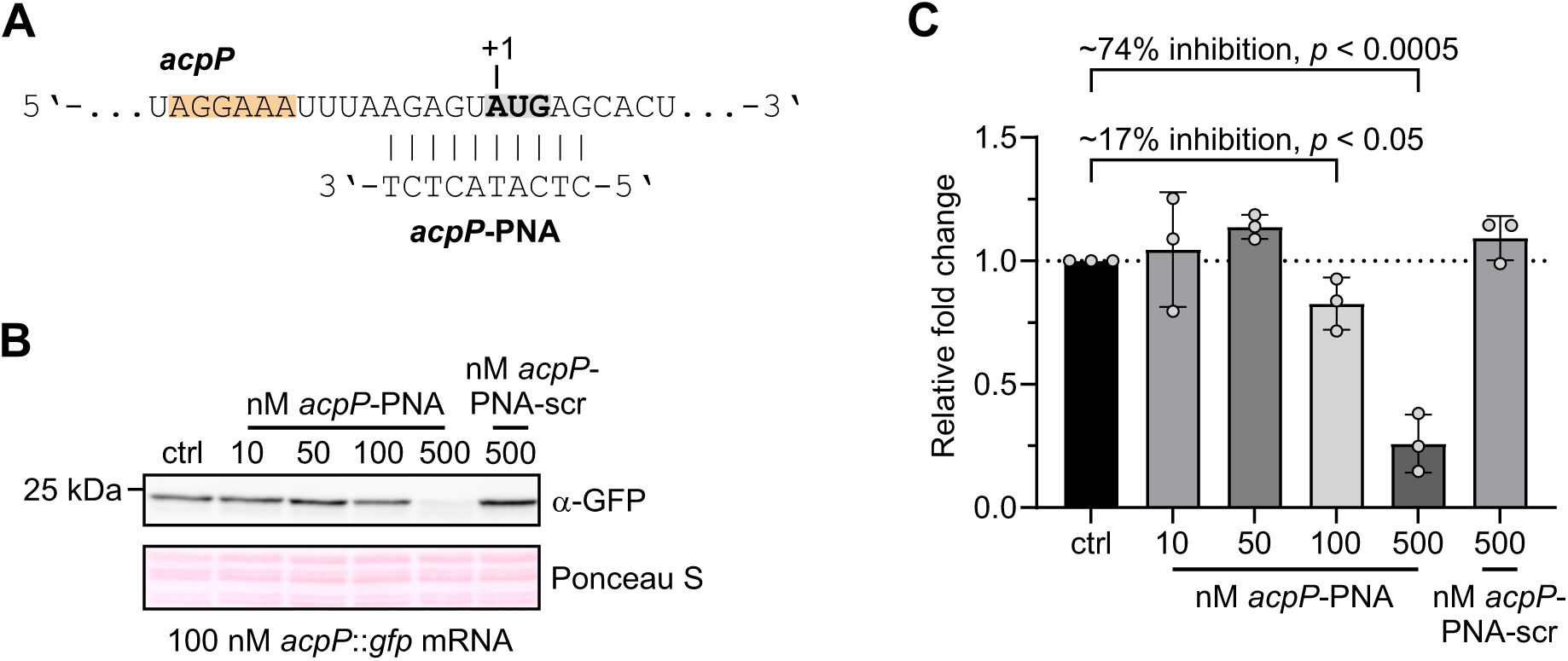
*In vitro* translational inhibition by antisense PNA. (A) Region of the *acpP* transcript targeted by *acpP*-PNA. Gray and bold, start codon. Orange, SD sequence. (B) Western blotting of an *in vitro* translation of an *acpP*::*gfp* fusion transcript reveals 608 reduced GFP protein levels with increasing concentrations of *acpP*-PNA. Ponceau S staining 609 was used as loading control. (C) Quantification of three independent western blots as shown in (B). Error bars show mean with SD. *p*-value was calculated using an unpaired, two-tailed t-test.

### Global analysis of PNA target inhibition using INRI-seq

Next, we set out to use INRI-seq to study the global effects of *acpP*-PNA on translation of its intended target as well as potential off-targets. We employed the same protocol that we used for the TIS analysis, except that we pre-annealed the synthetic transcriptome with varying concentrations of *acpP*-PNA or *acpP*-PNA-scr before starting the translation reactions. Similarly to the observed repression of the *acpP::gfp* fusion above (Figure 4B, C), INRI-seq reported a clearly reduced RPF density on the synthetic *acpP* mRNA with increasing *acpP*-PNA concentrations (Figure 5A). No inhibition of *acpP* translation was detected when *acpP*-PNA-scr was added, showing the specificity of PNA targeting. While the estimated concentration of *acpP* in our synthetic transcriptome was ∼500 pM (Supplementary Table S4), the actual concentration was likely higher: During the establishment of INRI-seq, we sequenced a test sample without addition of RET, revealing high levels of reads coming from the 3′ region of *acpP*, which was not part of the synthetic transcriptome (Figure S3A). This suggested that the *in vitro* translation kit probably contains traces of highly abundant RNAs, such as *acpP*. This hampers accurate estimations of ratios between *acpP*-PNA and its targets. Still, even low *acpP*-PNA concentrations showed a strong effect on *acpP* translation.

**Figure 5.**
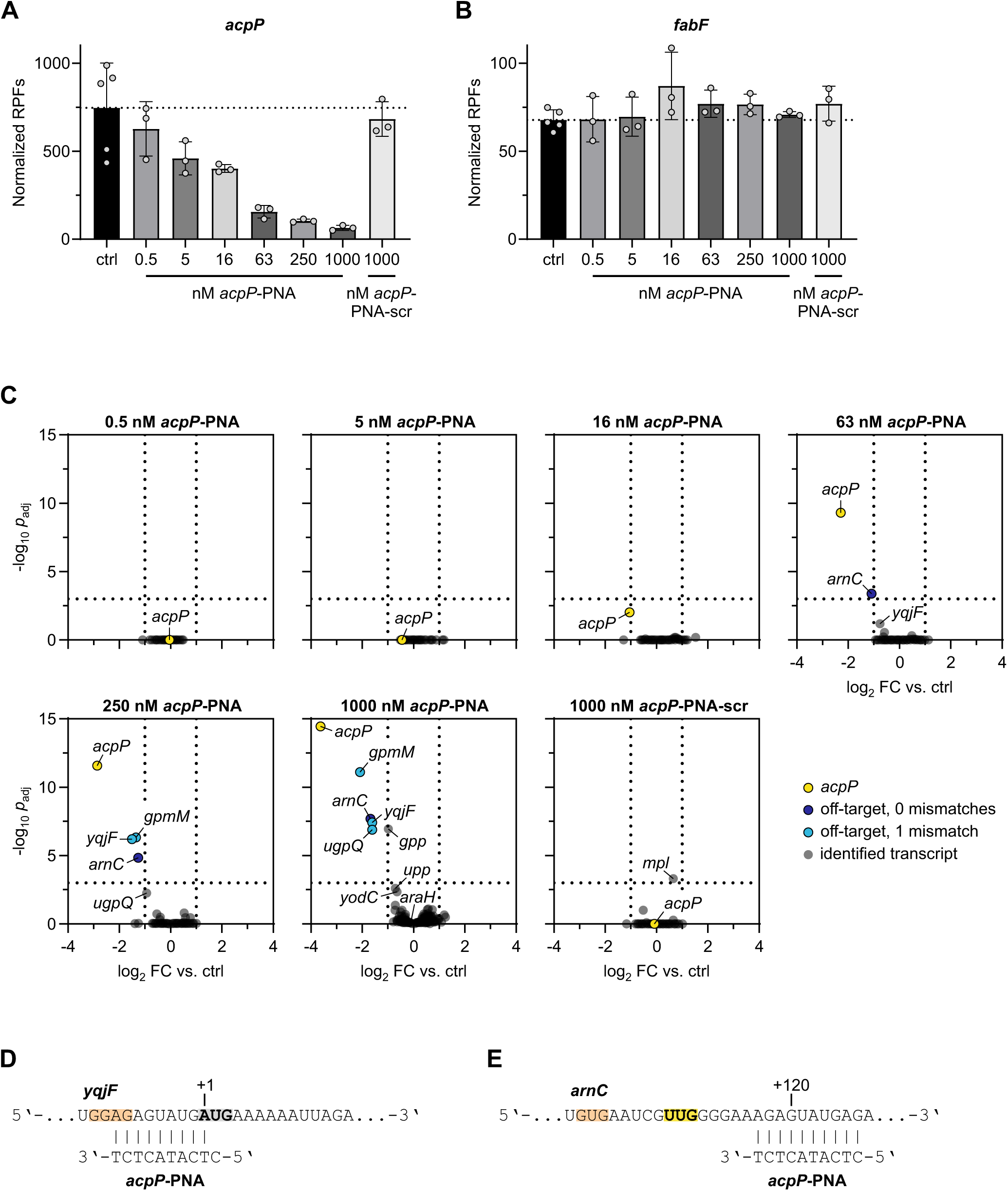
INRI-seq globally identifies targets of translation inhibition by PNA. (A) *acpP*-PNA inhibits translation of *acpP* in a concentration-dependent manner. Error bars show mean with SD. (B) *acpP*-PNA does not inhibit translation of *fabF*. Error bars show mean with SD. (C) Volcano plots showing calculated changes in translation as FDR-adjusted *p*-value against fold change. Cut-offs were set to *p*_adj_ < 0.001, log_2_ FC > |1| and are depicted as dotted lines. (D) Region of the *yqjF* transcript targeted by *acpP*-PNA. Gray and bold, start codon. Orange, SD sequence. (E) Region of the *arnC* transcript targeted by *acpP*-PNA. Yellow and bold, alternative start codon (compare to Figure S3E). Orange, SD sequence.

In the *E. coli* genome the *acpP* mRNA is co-transcribed with the downstream gene *fabF*^45^. *acpP*-PNA has been shown to lead to rapid decay of both cistrons following translation inhibition in cells, even though *fabF* is not a target of the antisense oligomer^28^. In agreement with the notion that *fabF* is down-regulated *in vivo* because its transcript is coupled to *acpP*, INRI-seq did not detect any change in translation of *fabF in vitro* (where the two transcripts are uncoupled), suggesting that INRI-seq faithfully reports only direct PNA-induced translational changes (Figure 5B).

### INRI-seq identifies direct PNA off-targets

We then analyzed the full spectrum of *acpP*-PNA-regulated targets. While *acpP* clearly was the primary transcript affected by the antisense oligomer, translation of a few other transcripts was also reduced at higher *acpP*-PNA concentrations (Figure 5C). One of them was *yqjF*, which has a stretch of 9 complementary bases to *acpP*-PNA within its translation initiation region (Figure 5D). Only the very 5′ nucleotide of the PNA is a mismatch (C:U) to *yqjF*. This suggests that at high concentration, *acpP*-PNA will also repress mRNAs that harbor a mismatch to the PNA sequence. Crucially, this was true for the additional off-targets *gpmM* and *ugpQ*. The translation initiation regions of both genes are complementary to *acpP*-PNA with only one mismatch at the 3′ end of the PNA (Figure 5C, Figure S3B, C). Finally, translation of the *gpp* transcript, which has complementarity to the antisense oligomer around its start codon, but with a T-G pair formed by the PNA’s second position (Figure S3D), was downregulated at 1 µM *acpP*-PNA as well, although it did not meet our strict cut-off conditions (*p*_adj_ < 0.001, log_2_ FC > |1|; Figure 5C).

The downregulated off-target *arnC* exhibits full complementarity to the antisense oligomer (Figure 5C, Supplementary Table S5). However, to our surprise, the *acpP*-PNA binds ∼120 nt downstream of the start codon of *arnC* (Figure 5E) where antisense repression of translation initiation is unlikely^41, 46, 47^. Interrogating our INRI-seq TIS dataset for alternative TISs within *arnC*, we indeed detected a peak in RPF density corresponding to an alternative in-frame UUG start codon, located a few nucleotides upstream of the *acpP*-PNA binding site (Figure S3E). Since no TISs were identified for *arnC* in the *in vivo* Ribo-RET data^18^, it remains unclear whether this alternative TIS is used *in vivo*.

Overall, these results suggest that, while target mismatches within the PNA sequence are generally thought to render the PNA inactive^42, 48^, mismatches at the 5′ or 3′ ends of the PNA are tolerated, leading to target regulation at high PNA concentration. In particular, of the 16 genes that harbor an *acpP*-PNA complementary sequence with one mismatch within their translation initiation region, six have the mismatch in position 1 or 10 of the PNA (Supplementary Table S5). Among those, the off-targets *gpmM, ugpQ* and *yqjF* are significantly downregulated, as discussed above. Of the other three, *upp* and *yodC* showed a tendency to be downregulated at the highest *acpP*-PNA concentrations, though they missed our statistical cut-offs (Figure 5C), while *araH* was not regulated at all. The ten genes, whose translation initiation regions include a complementary sequence to *acpP*-PNA with one mismatch in positions 2-8, showed no significant regulation in presence of the antisense oligomer. The same was true for all potential off-targets with two or more mismatches.

This analysis suggests that INRI-seq is able to accurately determine direct transcriptome-wide PNA-mediated inhibition of translation in response to defined PNA concentrations in a small reaction volume. Thus, it not only allows quantitative analysis of the direct effects of the PNA on the intended target transcript, but also enables the identification of PNA off-targets.

## DISCUSSION

In this study, we established INRI-seq, a cell-free method to globally study translation of a fully customizable synthetic transcriptome. Presenting two applications for this method, we identified known and new *E. coli* TISs and determined the fidelity of translation inhibition by antisense oligomers. Although optimized here for these purposes, the INRI-seq protocol is easily adaptable to other translation-related questions. For example, our protocol for PNA analysis can also be applied to the identification of the global targetome of sRNAs, including transcripts that have low abundance *in vivo*. This would allow the expansion of regulatory networks of known sRNAs and the investigation of the targets of currently understudied sRNAs. For single sRNA targets, *in vitro* translation has been successfully used to investigate sRNA-mediated regulation, indicating that a global study based on INRI-seq is feasible^47, 49, 50^. Importantly, INRI-Seq allows to include within the synthetic transcriptome targets that carry mutations in the putative sRNA binding site based on *in silico* predictions^51^. This enables testing specificity of targeting in the same experiment.

Since INRI-seq uses a fully synthetic transcriptome, users can analyze any desired sequence compilation. For example, the method can be used to compare translation of transcriptomes of related bacterial species or to analyze translation within complex microbial communities, such as the human gut microbiome. INRI-seq also enables the analysis of translation of phage transcripts independent of its host, facilitating the study of overlapping CDSs, which is a common feature in the complex genome organization of phages^52^, by disentangling them into single transcripts. Furthermore, mutational studies where a single or multiple transcripts are present in tens or even hundreds of different variants could be designed to investigate the influence of, for example, synonymous mutations on translation.

The design of the synthetic transcriptome is only limited by the length of the oligonucleotides: currently, up to 350 nt is commercially possible. In contrast to the length, the number of oligonucleotides in a pool (i.e., the available sequence space) generally is of less concern with several commercial providers, enabling the synthesis of more complex synthetic transcriptomes, including eukaryotic ones.

INRI-seq allows the study of translation in a quantitative manner, which is supported by the even coverage of transcripts in the synthetic transcriptome (Figure S1A) and read counts in the INRI-seq data (Figure S2B). Concentration-dependent analysis of translation-modulating molecules becomes feasible, because their exact concentration in the reaction is defined. This is not true in an *in vivo* setup, where the intracellular concentration is unknown. In addition, modulators can be added at high concentrations, which might otherwise lead to premature cell lysis when working with living cells. The small volume of the INRI-seq reactions (25 µl) is another advantage, especially if the translation modulator is limited or particularly expensive, like PNAs or antibiotic lead structures. Moreover, since there is no cell barrier or other resistance mechanisms to consider, comparative analysis of different types of antisense oligomers becomes possible, although there is no general mechanism for the cellular delivery of these compounds^24, 53, 54, 55x, 56^.

The commercial *in vitro* translation kit (PURExpress, New England Biolabs) used in the INRI-seq protocol is based on the PURE system^57^. It consists of individually purified *E. coli* components required for protein synthesis, such as initiation, elongation and termination factors as well as aminoacyl-tRNA synthetases, tRNAs, amino acids and ribosomes. This reaction mix allows efficient translation under most conditions. Nevertheless, the lack of factors like the translation elongation factor EF-P, which is necessary for efficient translation of polyproline stretches^58, 59^, could hamper the study of specific transcripts. In addition, the absence of most cellular RNA-binding proteins from of the kit and the shortening of the transcripts in the synthetic transcriptome may influence mRNA folding, which can impact translational efficiency^9^. RNA-binding proteins such as Hfq, which facilitates base pairing between sRNAs and their targets, are not part of the kit, either. Therefore, these proteins need to be purified and added to the reaction mixture when analyzing the effects of sRNAs, as previously done in an analysis of the sRNA GlmZ^50^. Finally, the presence of contaminating RNAs (Figure S1C), which are likely co-purified with the individual proteins of the kit, has to be kept in mind when designing concentration-sensitive, quantitative experiments. Despite these drawbacks, the PURE system works well for the applications tested here. By using other cell-free translation systems, such as self-generated S30 extracts from bacteria other than *E. coli*^60^ or those commercially available for human or rabbit^61^, we expect INRI-seq to be applicable to translational systems of all kingdoms of life.

In this study, we globally analyzed TISs of a synthetic *E. coli* transcriptome. Compared to a technically similar *in vivo* study^18^, we validated almost four times more annotated TISs (3,059 vs. 780), demonstrating the high sensitivity of INRI-seq. In addition, we detected 51 out of the 64 alternative TISs described by *in vivo* Ribo-RET^18^. INRI-seq further identified 918 additional putative alternative TISs for 858 genes, of which 279 are the only detected TISs for the respective genes (Figure 2D). These results are in agreement with a recent study suggesting that 5-12% of prokaryotic genes might have misannotated TISs^62^. Still, *in vivo* validation is required to verify whether these TISs are used within the cell.

To exploit the cell-free, low-volume nature of INRI-seq, we globally analyzed the influence of an antisense PNA on translation. In line with *in vivo* data^27, 28^, we found that *acpP*-PNA is specific for its target, *acpP*. Downregulation of PNA off-targets that harbor mismatches in the complementary sequence requires substantially higher PNA concentrations. In addition, mismatches are only tolerated in the terminal nucleotides of the PNA. Gratifyingly, a recent study investigating the effects of *acpP*-PNA on global transcript levels in a uropathogenic *E. coli* (UPEC) strain observed very similar off-target effects. Of the four off-targets identified by INRI-seq, *gpmM* and *ugpQ* were among the top regulated transcripts upon addition of *acpP*-PNA^27^. These findings should aid future PNA design by providing a framework to weigh the predicted specificity of PNAs according to their predicted off-targets.

Overall, INRI-seq provides a rapid, flexible *in vitro* protocol to investigate a wide range of translation-related questions. We expect this method to be an attractive alternative to *in vivo* Ribo-seq, particularly when availability or delivery of a translation-modulating molecule of interest is limiting.

## MATERIALS & METHODS

### Design and synthesis of the synthetic transcriptome

To obtain a synthetic version of the *E. coli* MG1655 transcriptome, a pool of single-stranded DNA oligonucleotides was designed by extracting the sequence of the first 153 nt of each annotated ORF (start codon + 50 codons). At the 5′ end, 30 nt upstream of each start codon was added as 5′ UTR and extended by addition of a T7 RNA polymerase promoter (GTTTTTTTTAATACGACTCACTATAGGG). At the 3′ end, a TAA stop codon was added and extended by the first 28 nt of the 3′ UTR of *E. coli hns* (TCTTTTGTAGATTGCACTTGCTTAAAAT). The final pool of 4,386 DNA oligonucleotides (JVOpool-001) was ordered from Integrated DNA Technologies at a scale of 1 pmol/oligo and is listed in Supplementary Table S6.

Using the KAPA HiFi HotStart PCR kit (Roche), double-stranded DNA was generated from JVOpool-001. Per 50 µl of PCR reaction, 20 ng of JVOpool-001, 10 µl HF buffer, 1 µl dNTPs, 1 µl DNA polymerase, and 2 µl each of the oligos JVO-18582 and JVO-18583 (10 µM each) were used. The PCR was run with the following protocol: 95°C for 3 min; 10 cycles of 95°C for 20 s followed by 60°C for 20 s and 72°C for 15 s; 72°C for 2 min. A total of 400 µl of this PCR was run and purified using column-based clean-up (Macherey-Nagel).

The RNA of the synthetic transcriptome was obtained by *in vitro* transcription of 500 ng each of the double-stranded DNA pool in two 40 µl reactions using the MEGAscript T7 transcription kit (Thermo Fisher) according to the manufacturer’s instructions, except that the reactions were performed overnight. The next day, the reactions were treated with 2 µl each of TURBO DNase (Thermo Fisher) at 37°C for 15 min. The RNA was denatured by addition of 40 µl of 2⨯ GLII loading buffer and incubation at 95°C for 5 min, then placed on ice. To purify the RNA, it was separated by 6% denaturing PAGE with 7 M urea in 1⨯ TBE, stained with EtBr, cut from the gel and eluted with 750 µl RNA elution buffer (0.1 M NaOAc, pH 5.4, 0.1 % SDS, 10 mM EDTA, pH 8) at 4°C overnight. The next day, the RNA-containing supernatant was mixed with 800 µl acidic phenol-chloroform-isoamylalcohol and centrifuged at 4°C for 15 min. The aqueous phase was collected, split into two tubes and precipitated with 1 ml each of ice-cold EtOH. Following precipitation at −20°C for at least 1 h, the samples were centrifuged at 4°C for at least 30 min, the supernatants discarded, the pellets washed with 400 µl ice-cold 70% EtOH and centrifuged again at 4°C for 15 min. Finally, the supernatants were discarded and the pellets were dried at room temperature for 5 min and resuspended in 25 µl water. The resuspended pellets were pooled into one tube to obtain the final synthetic transcriptome. RNA purity and integrity was tested by denaturing urea PAGE.

### In vitro Ribo-seq (INRI-seq)

To globally investigate translation *in vitro*, the PURExpress *In Vitro* Protein Synthesis Kit (New England Biolabs) was used to translate the synthetic transcriptome described above. To denature the synthetic transcriptome, it was incubated at 95°C for 2 min, then placed on ice. For each sample, a 25 µl reaction containing 10 µl solution A (PURExpress), 7.5 µl solution B (PURExpress), 5 µl water and 2.5 µl of the denatured synthetic transcriptome (stock concentration 10 µM; final concentration 1 µM) was incubated at 37°C for 15 min. Then, 1.25 µl of retapamulin (stock concentration 100 µg/ml in DMSO; final concentration 5 µg/ml) was added to block translation and the incubation continued at 37°C for 30 min. To stop the reaction, 175 µl of ice-cold stop buffer (20 mM Tris-HCl, pH 8, 100 mM NH_4_Cl, 10 mM MgCl_2_, 5 mM CaCl_2_, 1 mM DTT, 0.4 % Triton X-100, 0.1 % NP-40, 200 U/ml RNase-inhibitor, 5 µg/ml retapamulin) was added and the reaction put on ice.

When the effect of PNAs on translation was investigated, the same protocol was followed with the following exceptions: The PNA stock solutions (10⨯ the concentrations of the respective final concentrations) were incubated at 55°C for 5 min. Then, 2.5 µl of the PNA stock solution and 2.5 µl water were added to 2.5 µl of the denatured synthetic transcriptome and the mixture annealed at 37°C for 5 min, then placed on ice. The resulting 7.5 µl were then mixed with 10 µl solution A and 7.5 µl solution B and the translation carried out as described above.

RPFs were generated by addition of 1.4 µl of MNase (420 U in total; Thermo Fisher) and incubation at 25°C for 1 h. The reaction was quenched with 2 µl of 0.5 M EGTA, pH 8 and placed on ice. To isolate 70S monosomes containing the desired RPFs, 185 µl of the sample was loaded on a 10-55% sucrose gradient (20 mM Tris-HCl, pH 7.5, 100 mM NH_4_Cl, 10 mM MgCl_2_, 5 mM CaCl_2_, 1 mM DTT) formed in an open-top polyclear ultracentrifugation tube (Seton Scientific). The gradient was centrifuged at 4°C and 35,000 rpm for 2.5 h using an SW 40 Ti rotor (Beckman Coulter). Afterward, the 70S monosome fraction (∼1 ml) was collected using a Biocomp Model 153 gradient station and snap-frozen in liquid N_2_.

The frozen monosome fraction was thawed on ice and split into two tubes. To each tube, 800 µl of acidic phenol-chloroform-isoamylalcohol was added, the samples vortexed for 15 s and then centrifuged at 4°C for 15 min. The aqueous phases were collected and precipitated by addition of 1 µl GlycoBlue (Thermo Fisher) and 1.4 ml of ice-cold precipitation mix (30:1 EtOH:3 M NaOAc, pH 6.5) at −20°C for at least 1 h. The samples were centrifuged at 4°C for at least 30 min, the supernatants discarded, the pellets washed with 400 µl ice-cold 70% EtOH and centrifuged again at 4°C for 15 min. Finally, the supernatants were discarded and the pellets were dried at room temperature for 5 min and one pellet resuspended in 25 µl water. The solution was then transferred to the other tube to resuspend the corresponding second pellet.

30 µl of 2⨯ GLII loading buffer were added and the sample denatured at 95°C for 5 min, then placed on ice. As ladder, 10 µl of microRNA Marker (New England Biolabs) were mixed with 2.5 µl Low Range ssRNA Ladder (New England Biolabs), 25 µl 2⨯ GLII loading buffer and 22.5 µl water and then denatured like the sample. The RNA samples containing the RPFs and the ladder were separated by 15% denaturing PAGE with 7 M urea in 1⨯ TBE. The RPFs were visualized by staining the gel with SybrGold (Thermo Fisher) and cut from the gel in the range of 15-45 nt. Gel extraction was performed as described above, except that 1 µl GlycoBlue was added during ethanol precipitation. Finally, the pellets were dried at room temperature for 5 min and one pellet resuspended in 25 µl water. The solution was then transferred to the other tube to resuspend the corresponding second pellet.

### cDNA library preparation

The purified RPFs were subjected to library preparation for next-generation sequencing (vertis Biotechnologie). Briefly, the RNA 5′ ends were phosphorylated with T4 polynucleotide kinase (New England Biolabs). Adapters were ligated to the 5′ and 3′ ends of the RNA and first-strand cDNA synthesis was performed using M-MLV reverse transcriptase. The cDNA was then amplified using a high fidelity DNA polymerase and purified with Agencourt AMPure XP beads (Beckman Coulter). The cDNA samples were pooled in Equimolar amounts and the rDNA within the pool was depleted targeting 24 unique Sequences (Supplementary Table S1) using a CRISPR-Cas9 protocol similar to DASH^32^. The final pool was subjected to sequencing on an Illumina NextSeq 500 system using 75 nt single-end read length.

### Quantification of INRI-seq data

Reads of the INRI-seq experiments were preprocessed and mapped against the oligo pool using tools from the BBtools suite (https://sourceforge.net/projects/bbmap/). To remove sequencing adapters from raw reads and low-quality bases (Phred quality score < 10), BBduk was used. The resulting reads were mapped against the oligo pool using BBMap (v38.79). Aligned reads were then assigned genes and quantified using the featureCounts (v2.0.1) method of the Subread package^63^.

To facilitate the visualization of the 3′ ends of reads in coverage plots, wiggle files were created with the bamCoverage (v.3.4.3) tool from the deepTools package^64^. Here, the counts per million (CPM) normalization option was used to normalize for read depth per library. To get exact positional information on the ribosome position, only the last base of the 3′ end of each aligned read was profiled using the - Offset option of bamCoverage. The resulting coverages of the wiggle files were then visualized using integrative genomics viewer (IGV)^65^ and used for the identification of translation initiation sites.

### Analysis of translation inhibition by PNA

The R packages from the tidyverse-suite and edgeR (v3.30.0) were used to analyze the *in vitro* translation of the oligo pool^66, 67^. Briefly, a raw count table of quantified reads was imported into the edgeR environment. Oligos with < 4.21 CPM were removed. This cutoff was calculated by dividing 10/L, where L is the minimum library size in millions, as proposed by Chen *et al*.^68^. Next, read counts were normalized with edgeR’s trimmed mean of M values normalization method^69^. Differential translation was measured by first estimating the quasi-likelihood dispersions with the glmFit function and then comparing conditions with the glmQLFTest function. Transcripts with false discovery rate-adjusted^70^ *p*-values < 0.001 and log_2_ fold changes > 1 were considered to be differentially translated. In order to screen for possible off-targets of the *acpP*-PNA in the oligo-pool, the PNA sequence was mapped against the whole oligo pool using SeqMap, accepting alignments with up to one mismatch^71^.

### Metagene analysis

3′ end-aligned INRI-seq data was compared to an *in vivo* ribosome profiling dataset of *E. coli* pre-treated with retapamulin^18^. The raw coverage files were downloaded from GEO: GSE122129 and adjusted manually so that both datasets show the 3′ end of ribosomal footprints. To generate metagene plots, footprints from −10 to +55 relative to the start codon were extracted and the footprints were normalized against the total read depth of these reads. Then plots were created using functions of the R packages ggplot2 and the dplyr^67^.

### Computational analysis of translation initiation sites

To identify annotated TISs, an algorithm was implemented using custom python scripts. Briefly, the algorithm scans through each of the oligo sequences base by base and looks for peaks. For annotated TISs, it searches for peaks in the region of 15 (± 3) bases inside the CDS, because that is where the 3′ end of the reads is located during translation initiation, when a ribosome is attached^31^. If a peak was identified (CPM normalized depth > 5) in at least 2 replicates, it was considered a TIS in the INRI-seq dataset (Figure S4A, Supplementary Table S2).

In addition to the confirmation of annotated TISs, *in vivo* retapamulin-treated ribosome profiling had been used previously to search for alternative TISs, which do not coincide with annotated start codons^18^. Custom python scripts were run to find alternative TISs and to compare them to the *in vivo* data (Figure S4B). Each oligo was screened for peaks with > 5 CPM outside its annotated TIS. Only peaks with a relative density (reads of peak divided by the total reads for the respective oligo) > 10% were considered. Additionally, only peaks with a start codon (AUG, UUG, GUG, CUG, AUC, and AUU) 15 nt (± 3nt) upstream of the peak were considered. To prevent the identification of more than one alternative TIS for the same site, peaks in close proximity (up to 5 nt in distance) were merged and the highest respective peak was selected. Next, the reading frame of the alternative TIS relative to the annotated reading frame was noted and a stop codon downstream of the alternative TIS was searched for, designating the alternative TIS as in-frame or out-of-frame. Finally, the *in vivo* Ribo-RET data^18^ was searched for the alternative TIS using the same settings.

### In vitro translation followed by western blotting

The *acpP*::*gfp* fusion transcript was obtained as previously described^72^. Briefly, *E. coli* genomic DNA was PCR amplified with JVO-18305 and JVO-18306 (Supplementary Table S7) and cut with NheI and NsiI. Plasmid pXG-10^72^ was also cut with NheI and NsiI and ligated with the *acpP* insert. After transformation and plasmid isolation, the plasmid containing the *acpP*::*gfp* fusion template was PCR amplified with JVO-18200 and revSF to obtain the dsDNA for *in vitro* transcription. After *in vitro* transcription and purification as described above, *in vitro* translation of the *acpP*::*gfp* fusion transcript was carried out as for INRI-seq with the following exceptions: Translation reaction volumes were scaled down to a total of 10 µl, the final transcript concentration was 100 nM, no RET was added to the reactions and translation was carried out for 2 h. After the incubation, 2.5 µl of 5⨯ protein loading buffer was added and the samples incubated at 95°C for 2 min and placed at room temperature. The samples were then separated by 12% SDS-PAGE followed by blotting onto a PVDF membrane (GE Healthcare). The membrane was stained with Ponceau S (Sigma-Aldrich) as loading control and incubated with a mouse α-GFP antibody (Roche). An HRP-coupled goat α-mouse antibody (Thermo Fisher) was finally used to develop the blot.

## DATA AVAILABILITY

The sequencing data have been deposited in NCBI’s Gene Expression Omnibus^73^ and are accessible through GEO Series accession number GSE190954 (https://www.ncbi.nlm.nih.gov/geo/query/acc.cgi?acc=GSE190954). The custom code used to analyze the sequencing data can be found on GitHub (https://github.com/BarquistLab/oligopool_riboseq).

## ACKNOWLEDGMENTS

We thank the members of the Vogel lab for splendid ideas and discussions, Lydia Hadjeras and Sarah Svensson for their invaluable help to set up ribosome profiling, and Anke Sparmann for editing the manuscript. We thank the Core Unit SysMed at the University of Würzburg for excellent technical support and RNA-seq data generation. Research was supported by the Bavarian Bayresq.net project Rbiotics (LB, JV) and a German Research Council Leibniz Award (JV, grant number DFG Vo875-18).

## AUTHOR CONTRIBUTIONS

Conceptualization: JH, JV. Methodology: JH, JJ, LB, JV. Software: JJ, LB. Validation: JH, JJ, SD-M. Formal analysis: JH, JJ. Investigation: JH. Resources: LB, JV. Data Curation: JJ, LB. Writing – Original Draft: JH, JV. Writing – Review & Editing: JH, JJ, SD-M, LB, JV. Visualization: JH. Supervision: LB, JV. Project Administration: LB, JV. Funding acquisition: LB, JV.

## CONFLICT OF INTEREST STATEMENT

The authors declare no competing interests.

## Notes

### Competing Interest Statement

The authors have declared no competing interest.

