## Supplementary Figures for "INRI-seq enables global cell-free analysis of translation initiation and off-target effects of antisense inhibitors"

#### SUPPLEMENTARY FIGURE LEGENDS

**Figure S1.** Quality control of INRI-seq.

(A) Violin plot showing the read distribution of all transcripts in the synthetic transcriptome with > 10 reads per million. The dashed line represents the median and the dotted lines represent the 25% and 75% percentiles.

(B) Sucrose gradient UV profiles of the sedimentation of *in vitro* translation reactions with or without addition of RET.

(C) Read distribution of rRNA genes without rRNA depletion.

(D) Comparison of the read distribution of RNA classes with and without rRNA depletion.

**Figure S2.** Reproducibility of INRI-seq.

(A) Scatter plot showing the correlation between two representative replicates. $R^2$ represents the coefficient of determination.

(B) Violin plots showing the read coverage of the 4,149 transcripts detected in all five replicates (R1-R5). The dashed lines represent the medians and the dotted lines represent the 25% and 75% percentiles.

(C) The average distance of RPF peak density from annotated start codons is 15 nt for each of the replicates, and their RPF densities are congruent.

(D, E) RPF density at the TIS of *hfq*.

**Figure S3.** Evaluation of PNA off-targets.

(A) RPF distribution of *acpP* without addition of RET. The part of *acpP* included in the synthetic transcriptome is indicated.

(B-D) Regions of the *gpmM*, *ugpQ* and *gpp* transcripts targeted by *acpP*-PNA. Gray and bold, start codon. Orange, SD sequence.

(E) INRI-seq identifies an RPF peak within *arnC* belonging to an in-frame alternative start codon (yellow and bold). Orange, SD sequence.

**Figure S4.** Overview of TIS analysis.

(A) Analysis of annotated TISs.

(B) Analysis of putative new TISs.

### Figure S1

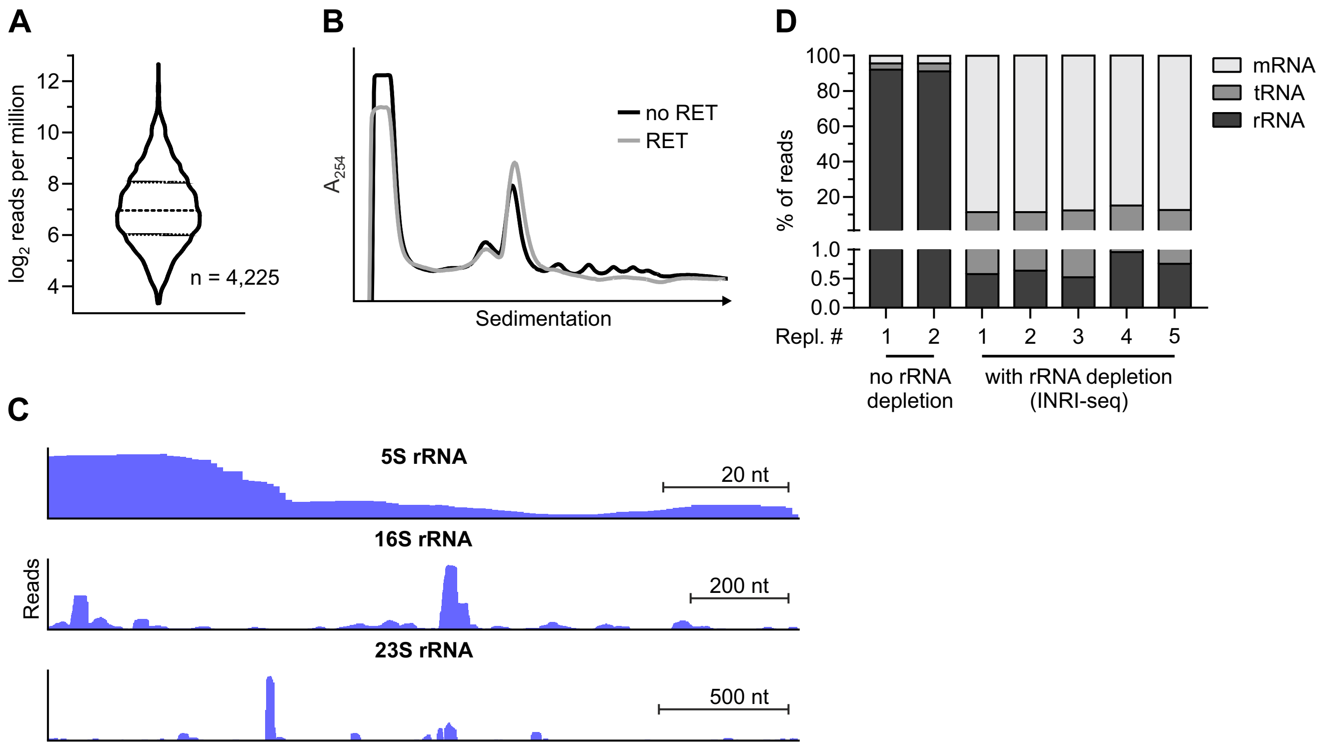

A
log2 reads per million
12
10
8
6
4
n = 4,225
B
A254
no RET
RET
Sedimentation
C
5S rRNA
20 nt
16S rRNA
Reads
200 nt
23S rRNA
500 nt
D
100
80
60
40
20
% of reads
1.0
0.5
0.0
mRNA
tRNA
rRNA
Repl. #
1
2
1
2
3
4
5
no rRNA depletion
with rRNA depletion (INRI-seq)

### Figure S2

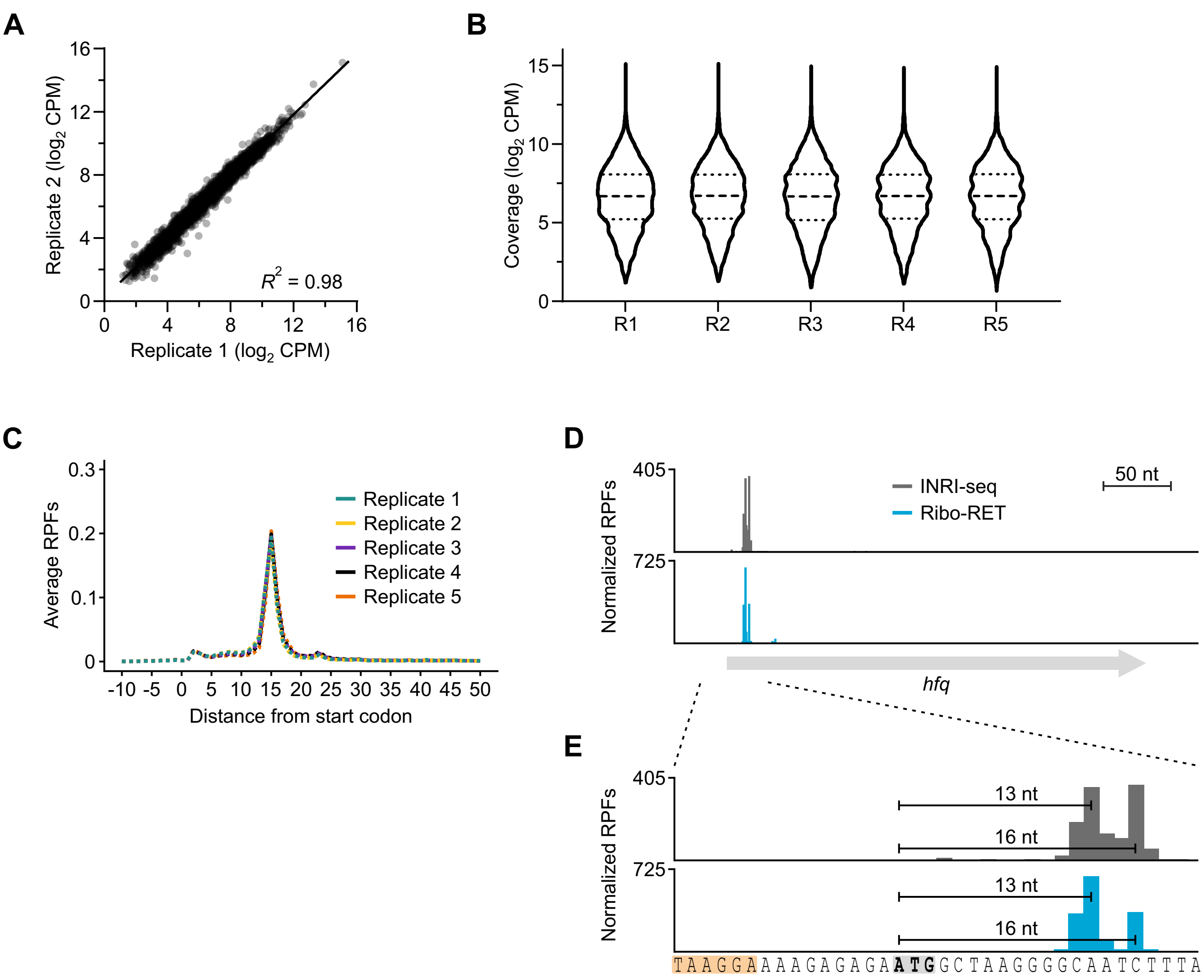
A
Replicate 2 (log2 CPM)
Replicate 1 (log2 CPM)
R2 = 0.98
B
Coverage (log2 CPM)
R1
R2
R3
R4
R5
C
Average RPFs
Replicate 1
Replicate 2
Replicate 3
Replicate 4
Replicate 5
Distance from start codon
D
Normalized RPFs
405
725
INRI-seq
Ribo-RET
50 nt
hfq
E
Normalized RPFs
405
725
13 nt
16 nt
13 nt
16 nt
TAAGGAAAAGAGAGAATGGCTAAGGGGCAATCTTTA

### Figure S3

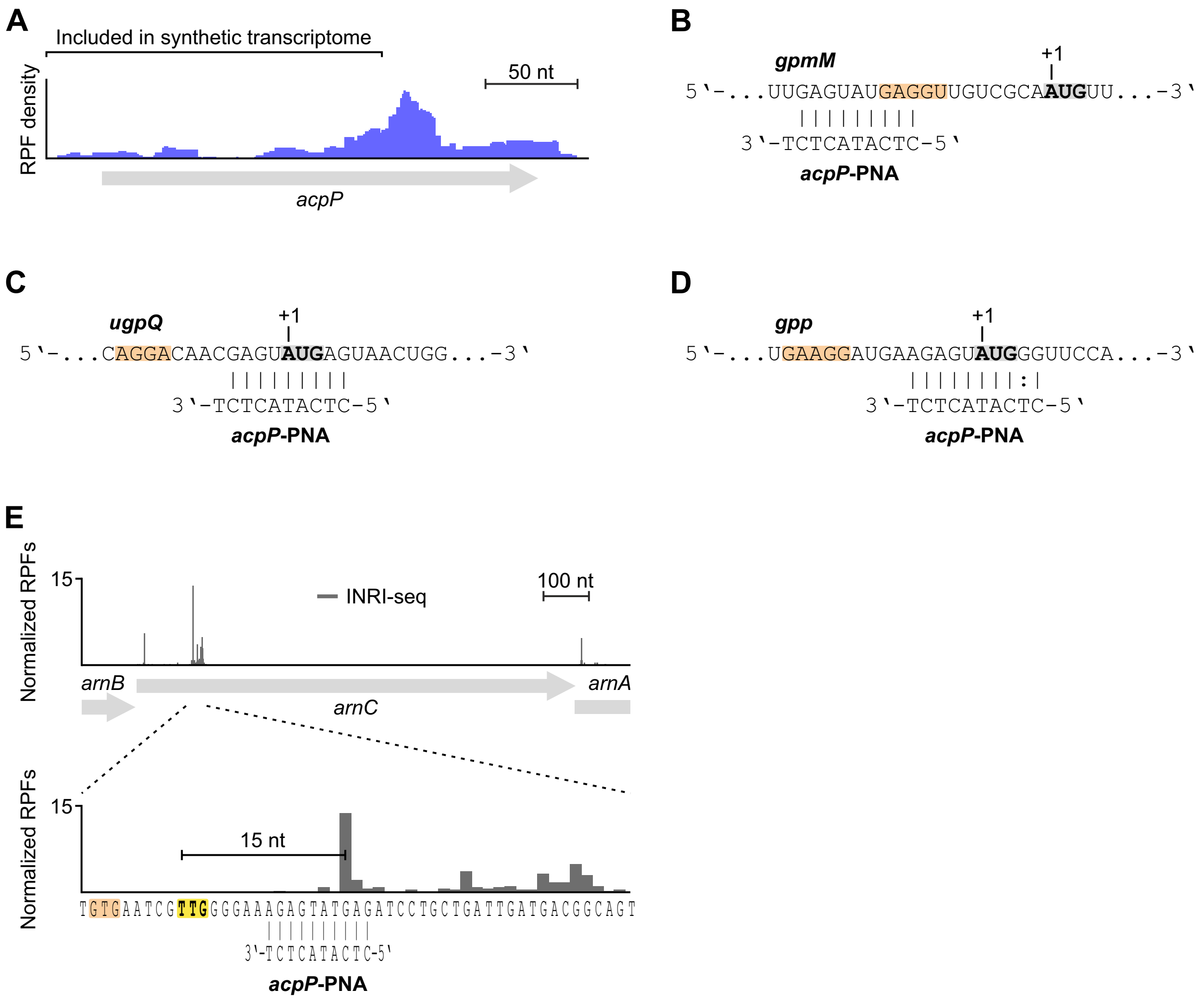

A
Included in synthetic transcriptome
50 nt
RPF density
acpP
B
gpmM
+1
5'-...UUGAGUAUGAGGUUGUCGCAAUGUU...-3'
3'-TCTCATACTC-5'
acpP-PNA
C
ugpQ
+1
5'-...CAGGACAACGAGUAUGAGUAACUGG...-3'
3'-TCTCATACTC-5'
acpP-PNA
D
gpp
+1
5'-...UGAAGGAUGAAGAGUAUGGGUUCCA...-3'
3'-TCTCATACTC-5'
acpP-PNA
E
Normalized RPFs
15
INRI-seq
100 nt
arnB
arnC
arnA
Normalized RPFs
15
15 nt
TGTGAATCGTTGGGGAAAGAGTATGAGATCCTGCTGATTGATGACGGCAGT
3'-TCTCATACTC-5'
acpP-PNA

### Figure S4

A

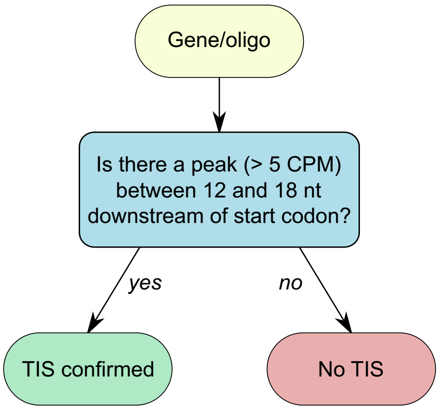
Gene/oligo
Is there a peak (> 5 CPM) between 12 and 18 nt downstream of start codon?
yes
no
TIS confirmed
No TIS

B

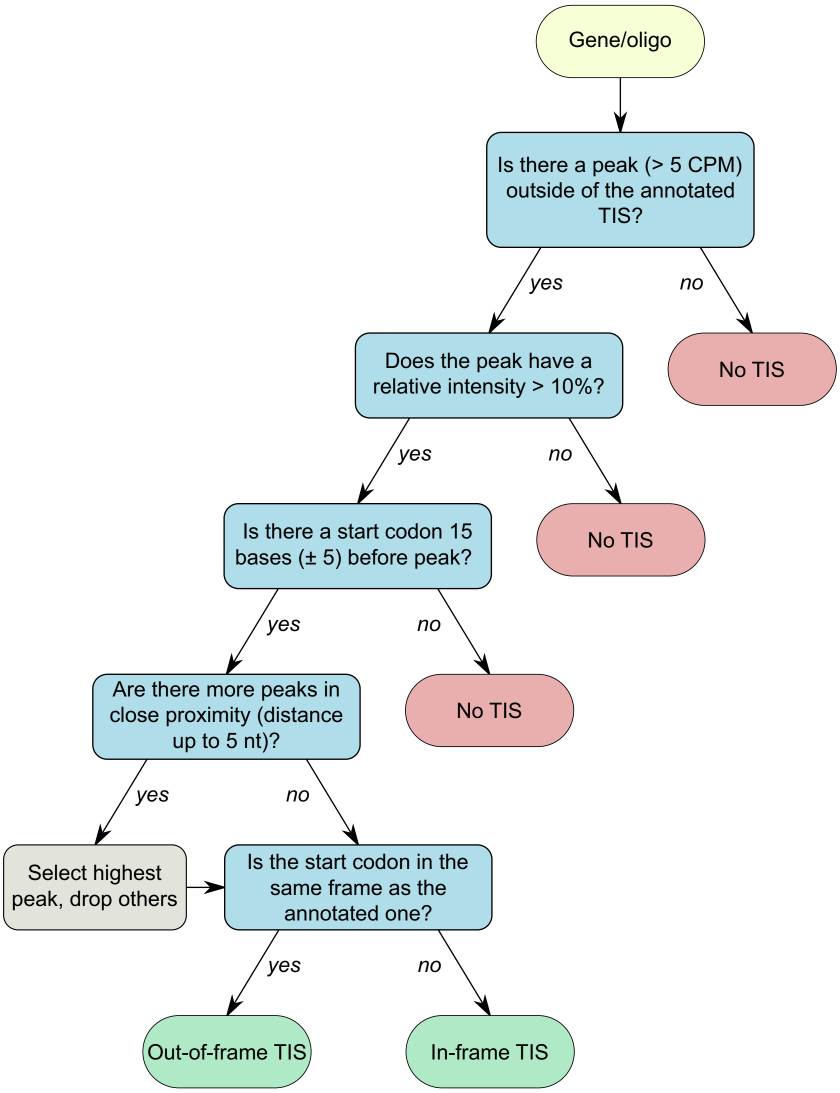
Gene/oligo
Is there a peak (> 5 CPM) outside of the annotated TIS?
yes
no
No TIS
Does the peak have a relative intensity > 10%?
yes
no
No TIS
Is there a start codon 15 bases (± 5) before peak?
yes
no
No TIS
Are there more peaks in close proximity (distance up to 5 nt)?
yes
no
Select highest peak, drop others
Is the start codon in the same frame as the annotated one?
yes
no
Out-of-frame TIS
In-frame TIS
